# Hypocretin receptor 1 blockade early in abstinence reduces future demand for cocaine

**DOI:** 10.1101/2024.12.06.627226

**Authors:** Shanna B. Samels, Jessica K. Shaw, Pamela Alonso, Emily M. Black, Rodrigo A. España

**Affiliations:** Department of Neurobiology and Anatomy, Drexel University College of Medicine, Philadelphia, PA 19129; Department of Psychiatry, University of Pennsylvania, Philadelphia, PA 19104; Department of Physiology, Louisiana State University, New Orleans, LA 70112; Department of Psychology, Haverford College, Haverford, PA 19041

## Abstract

Relapse to cocaine use after abstinence remains a significant challenge for treating cocaine use disorder. While the mechanisms of relapse are still under investigation, adaptations in mesolimbic dopamine systems may contribute to cocaine craving and propensity for relapse. Current pharmacological treatments targeting dopamine systems are often intolerable and may have abuse potential. Therefore, identifying novel pharmacological targets for cocaine use disorder is crucial. The hypocretin/orexin system has been shown to regulate cocaine-associated behavior and dopamine transmission. Our previous studies indicated that the hypocretin receptor 1 antagonist, RTIOX-276, reduced motivation for cocaine and attenuated dopamine responses to cocaine. Importantly, the effects of RTIOX-276 on dopamine transmission persisted for at least 24 hours, suggesting lasting effects of hypocretin receptor antagonism. Here, we hypothesized that a single RTIOX-276 treatment would reduce motivation for cocaine and normalize dopamine transmission after abstinence. Rats were pre-assessed for cocaine consumption and motivation using a within-session threshold schedule before intermittent access exposure to cocaine. Rats were subsequently treated with RTIOX-276 on the first day of a 7-day abstinence period, after which they were reassessed for cocaine consumption and motivation or examined for dopamine transmission using fast-scan cyclic voltammetry in nucleus accumbens core slices. We found that a single treatment with RTIOX-276 on the first day of abstinence reduced motivation for cocaine and normalized aberrant dopamine uptake observed following intermittent access to cocaine. These findings suggest that hypocretin receptor 1 may be a viable target for reducing motivation for cocaine through alterations in dopamine transmission in the nucleus accumbens.

## Introduction

Abstinence from drug use has been shown to elicit a progressive intensification or ‘incubation’ of drug craving/seeking (Bedi et al., 2011; Wang et al., 2013; Li et al., 2015) that is posited to underlie the risk of relapse (Grimm et al., 2001; Grimm et al., 2003; Bienkowski et al., 2004; Mead et al., 2007; Abdolahi et al., 2010). Accumulating evidence suggests that mesolimbic dopamine adaptations develop during cocaine abstinence and may contribute to incubation cocaine seeking (Calipari et al., 2013; Calipari et al., 2015; Wolf, 2016; Alonso et al., 2021; Samaha et al., 2021). Unfortunately, pharmacotherapies that target dopamine systems directly are largely ineffective or intolerable and may have abuse potential themselves. Therefore, identifying novel treatments for cocaine use disorder is crucial.

The hypocretin/orexin (Hcrt) neuropeptide system has been demonstrated to regulate dopamine transmission and influence motivation for cocaine (Shaw et al., 2019; Brodnik et al., 2020). For example, antagonism of Hcrt receptor 1 (Hcrtr1) reduces high effort self-administration of cocaine and disrupts dopamine transmission at baseline and following exposure to cocaine (Borgland et al., 2009; España et al., 2010; Brodnik et al., 2015; Prince et al., 2015; Levy et al., 2017). Importantly, these prior studies assessed the acute, on-board pharmacological effects of Hcrtr1 antagonism on behavior. However, we and others have reported that the effects of Hcrtr1 antagonists may be long lasting – extending beyond the on-board, pharmacological effects (Ishii et al., 2005; Winrow et al., 2010; Brodnik et al., 2020; Mohammadkhani et al., 2020). For instance, the Hcrtr1 antagonist RTIOX-276 produced lasting effects on dopamine transmission that were maintained for at least 24 hours after administration (Brodnik et al., 2020). Moreover, in a recent study we showed that a single treatment with RTIOX-276 early in cocaine abstinence prevented incubation of cocaine seeking 7 days later. However, in that study rats were tested for cue-induced cocaine seeking, during which rats do not receive cocaine (Clark et al., 2024). Consequently, it remains unclear whether HcrtR1 antagonism early in abstinence reduces active cocaine self-administration later in abstinence.

In the present studies, we tested the lasting effects of a single RTIOX-276 treatment early in cocaine abstinence on later motivation for cocaine and to what extent these effects were associated with alterations in dopamine transmission. Rats initially self-administered using a within-session threshold schedule of reinforcement to obtain a pre-assessment of cocaine consumption and motivation (Oleson and Roberts, 2009; España et al., 2010; Oleson et al., 2011; Oleson and Roberts, 2012; Bentzley et al., 2013; Brodnik et al., 2017; Shaw et al., 2021).To promote incubation of cocaine seeking and enhance dopamine responses to cocaine, rats were then exposed to 7 days of intermittent access (IntA) (Calipari et al., 2013; Calipari et al., 2015; Alonso et al., 2022; Clark et al., 2024). On the first day of a 7-day abstinence period (abstinence day 1; AD1) rats were treated with either intravenous vehicle or 20mg/kg RTIOX-276. To determine the lasting effects of RTIOX-276 on behavior, one group of rats underwent a post-assessment using the threshold schedule on abstinence day 8 (AD8). A separate group of rats underwent the same behavioral conditions prior to abstinence but were prepared for fast scan cyclic voltammetry (FSCV) on AD8 instead of being tested on the threshold schedule. Together these experiments allowed us to test the impact of Hcrt antagonist pretreatment on motivation for cocaine after abstinence, and to identify changes in dopamine transmission that contribute to observed behavioral changes.

## Materials and Method

### Animals

Adult female (200g -250g) and male (300g -350g) Sprague Dawley rats (Envigo, Frederick, MD, USA) were maintained on a 12-hour reverse light/dark cycle (1500 lights on; 0300 lights off) with *ad libitum* access to food and water. After arrival, rats were given 7 days to acclimate prior to surgery. All protocols and animal procedures were conducted in accordance with the National Institute of Health Guide for the Care and Use of Laboratory Animals under supervision of the Institutional Animal Care and Use committee at Drexel University College of Medicine.

### Drugs

Cocaine hydrochloride was provided by the National Institute on Drug Abuse, Drug Supply Program (Research Triangle Park, NC, USA). For self-administration experiments, cocaine was dissolved in 0.9% physiological saline. For FSCV experiments, cocaine was dissolved in artificial cerebrospinal fluid (aCSF). RTIOX-276 was provided by Drs. Zhang and Perrey (Research Triangle Institute) (Perrey et al., 2013). The dose of RTIOX-276 used was 20 mg/kg (Perrey et al., 2013; Levy et al., 2017). RTIOX-276 was dissolved in 5% Tween20 and pH was adjusted to ∼7.5 with 1M NaOH and 1M HCL solutions.

### Intravenous catheter surgery

Rats were anesthetized using 2.5% isoflurane and implanted with a silastic catheter inner diameter (ID) of 0.012 in., outer diameter (OD) of 0.025 in. (Access Technologies, Skokie, IL) into the right jugular vein for intravenous delivery of cocaine. The catheter was connected to a cannula that excited the skin on the dorsal surface in the region of the scapula. Ketoprofen (Patterson Veterinary, Devens, MA; 5, mg/kg s.c of 5mg/ml) and Enrofloxacin (Norbrook, Northern Ireland; 5, mg/kg s.c of 5mg/ml) were given at the time of surgery and a second dose was given 12 hr later. Antibiotic/analgesic powder (Neopredef, Kalamazoo, MI) was applied around the chest and back incisions. Rats were then singly housed and allowed to recover for 3 days prior to self-administration training. Intravenous catheters were manually flushed with heparinized saline during recovery to maintain catheter patency.

### Self-administration

All measures self-administration measures were recorded using custom created Ghost Software (Bernosky-Smith et al., 2016). Rats underwent one cocaine self-administration session per day from 10:00-16:00. Rats were first trained to self-administer cocaine on a 6-hr fixed ratio 1 (FR1) schedule where a single active lever response initiated an intravenous injection of cocaine (0.75 mg/kg, infused over 2.5 – 4s) paired with a cue light above the active lever. Each rewarded lever press was followed by a 20s timeout where the lever retracted. Rats acquired cocaine self-administration when they reached 20 infusions in one 6-hr FR1 session. Rats were then switched to a 2-hr FR1 schedule to maintain self-administration before being switched to within-session threshold by achieving 20 infusions per session across three consecutive sessions.

### Pre-Assessment Within-Session Threshold

After maintenance of FR1 self-administration, rats underwent a 3-day pre-assessment for consumption and motivation for cocaine using the within-session threshold procedure. Under this schedule, the dose of cocaine is initially high and then decreases in a stepwise, quarter-logarithmic fashion every 10 min for 110 min (435.7, 245, 137.8, 77.5, 43.6, 24.5, 13.8, 7.7, 4.4, 2.4 1.4 µg/infusion) (España et al., 2010; Oleson and Roberts, 2012). At the beginning of the session the dose of cocaine is relatively high, and rats can titrate their preferred blood level of cocaine with minimal effort. This procedure provides measures of consumption (intake when cocaine is ‘essential free’, Q_0_), intake (mg per the 2-3 10 min epochs), demand elasticity (α; an inverse measure of motivation) and maximal effort expended to maintain preferred levels of cocaine (Pmax). The values of consumption (Q_0_) and demand elasticity (α) were calculated using an automated script in RStudio that generated cocaine demand curves that fit the least squares regression curve to the natural log (mg intake) by unit price of cocaine at each epoch (Oleson et al., 2011). Consumption (Q_0_) was defined as the y-intercept and demand elasticity (α) was defined as the slope of the curve (Hursh and Silberberg, 2008). Intake was calculated by averaging the intake across three epochs in the early phase when rats are titrating preferred cocaine levels (España et al., 2011). The first “epoch” was excluded from the analysis, given that prior work demonstrated that during the first epoch rats ‘load up’ on cocaine to quickly achieve high blood levels of cocaine (España et al., 2010; Oleson et al., 2011; Bentzley et al., 2013). Pmax was calculated as the unit price at which maximal responding occurred (España et al., 2010; España et al., 2011). These values were averaged across 3 sessions of the within-session threshold test for each rat.

### Intermittent Access (IntA)

Following the pre-assessment on the within-session threshold procedure, rats were switched to the IntA schedule of reinforcement. On this schedule rats had access to cocaine for 5 min followed by a 25 min timeout during which levers retract. This 30-min trial was repeated for 12 trials during a 6-hr session each day. Lever presses resulted in a single intravenous infusion of cocaine (0.375 mg/kg), paired with a cue light above the lever. During the 5 min periods when the lever was available there was no timeout following lever presses. Rats underwent IntA for 7 consecutive days before moving on to abstinence.

### Abstinence

Following the final IntA self-administration session, rats underwent a forced abstinence period for 7 days. During this time, rats remained in their testing chambers, but no self-administration sessions occurred. On AD1, rats received either vehicle or 20mg/kg RTIOX-276. Rats were flushed with heparinized saline on abstinence day 2 through 7 to maintain catheter patency.

### Post-Assessment Within-Session Threshold

Following IntA and abstinence, rats were reassessed for consumption (Q_0_), intake, demand elasticity (α), and Pmax using additional within session threshold sessions for 3 days. These post-assessment threshold measures were averaged across the 3 sessions for each rat and were expressed as both pre-vs. post-assessment raw values and as a percent of the pre-assessment threshold averages.

### Fast scan cyclic voltammetry (FSCV)

Following IntA and abstinence a separate group of rats was euthanized on AD8 for FSCV in brain slices. These rats did not undergo post-assessment threshold tests to avoid the confounding influences of re-exposure to cocaine, which has been shown to induce rapid alterations in dopamine transmission following abstinence (Siciliano et al., 2016). Rats were anesthetized with 2.5% isoflurane for 5 min and decapitated. The brain was rapidly dissected and transferred to ice-cold aCSF containing NaCl (126 mM), KCl (2.5 mM), NaH2PO_4_ (1.2mM), CaCl_2_ (2.4 mM), MgCl_2_ (1.2 mM), NaHCO_3_ (25 mM), glucose (11 mM), and L-ascorbic acid (0.4 mM), pH adjusted to 7.4. A vibratome was used to slice 400 µM thick coronal sections containing the NAc core. Slices were transferred to room temperature oxygenated aCSF and left to equilibrate for at least 1 hr before being transferred into a recording chamber with aCSF (32°C). Cocaine-naive (naive) rats were used as a control group throughout these studies. A bipolar stimulating electrode was placed on the surface of the tissue in the NAc core, and a carbon fiber microelectrode was implanted between the stimulating electrode leads. dopamine release was evoked every 3 min using a single electrical pulse (400 µA, 4ms, monophasic) and measured using Demon Voltammetry and Analysis Software (Yorgason et al., 2011). Following collections of baseline dopamine release (3 consecutive stimulations within <10% variation), the slice was exposed to increasing cocaine concentrations (0.3, 1.0, 3.0, 10, 30µM). FSCV data were analyzed using Demon Voltammetry and Analysis Software using Michaelis-Menten kinetic methods to calculate dopamine release, maximal rate of dopamine uptake, and dopamine transporter (DAT) sensitivity to cocaine (i.e., cocaine-induced inhibition of dopamine uptake; apparent K_m_) (Yorgason et al., 2011).

### Data analysis and statistics

Statistical analyses were conducted using GraphPad Prism 10 except three-way ANOVAs were performed using SPSS Statistics (Version 29). Analyses to detect sex differences were conducted for all behavioral and voltammetry metrics using either a two-way ANOVA or a three-way mixed design ANOVA. We did not observe interactions between sex and RTIOX-276 treatment for any measure of interest indicating that both females and males responded similarly across test conditions. Therefore, female and male data were combined. Table S1 shows results from the sex differences analyses performed. Behavioral data were analyzed using Student’s t-tests or two-way ANOVAs. Baseline voltammetry measures of release and uptake were analyzed using one-way ANOVAs. The effects of cocaine on release and inhibition of uptake were analyzed using two-way ANOVAs. Specific analyses are detailed in the results.

## Results

### Blockade of Hcrtr1 early in abstinence reduces motivation for cocaine later in abstinence

To assess to what extent HcrtR1 antagonism early in abstinence reduces motivation for cocaine after 7 days of abstinence, rats were first tested on the threshold schedule to obtain a pre-assessment of cocaine consumption and motivation prior to self-administering on IntA for 7 days and prior to receiving vehicle (n=10) or 20 mg/kg RTIOX-276 (n=8) treatment on AD1 (**Figure 1A**). Student’s t-tests revealed no differences in days to acquire cocaine self-administration (t_(36)_=0.6704, *p*=0.5069; Figure 1B), consumption (t_(36)_=0.4013, *p*=0.6905; **Figure 1C**), intake (t_(36)_=0.4013, *p*=0.6905; **Figure 1D**), demand elasticity (t_(36)_=0.7879, *p*=0.9376; **Figure 1E**) or Pmax (t_(36)_=0.9613, *p*=0.3428; **Figure 1F**) between rats that would eventually receive vehicle (Pre-Veh) or RTIOX-276 (Pre-RTI). Similarly, a two-way ANOVA with future RTIOX-276 treatment (Pre-Veh vs Pre-RTI) as the between subjects variable and day of self-administration (days 1-7) as the within-subjects variable revealed no effect of future RTIOX-276 treatment (*F*_(1,36)_=0.09092, *p*=0.7647), day (*F*_(3.391,122.1)_=0.7555, *p*=0.5361) nor future RTIOX-276 treatment x day interaction (*F*_(6,216)_=0.5924, *p*=0.7363) on the number of infusions taken over the course of IntA self-administration (**Figure 1G**). Together these observations indicate there were no differences in self-administration behavior between groups that would eventually receive vehicle vs RTIOX-276.

**Figure 1:**
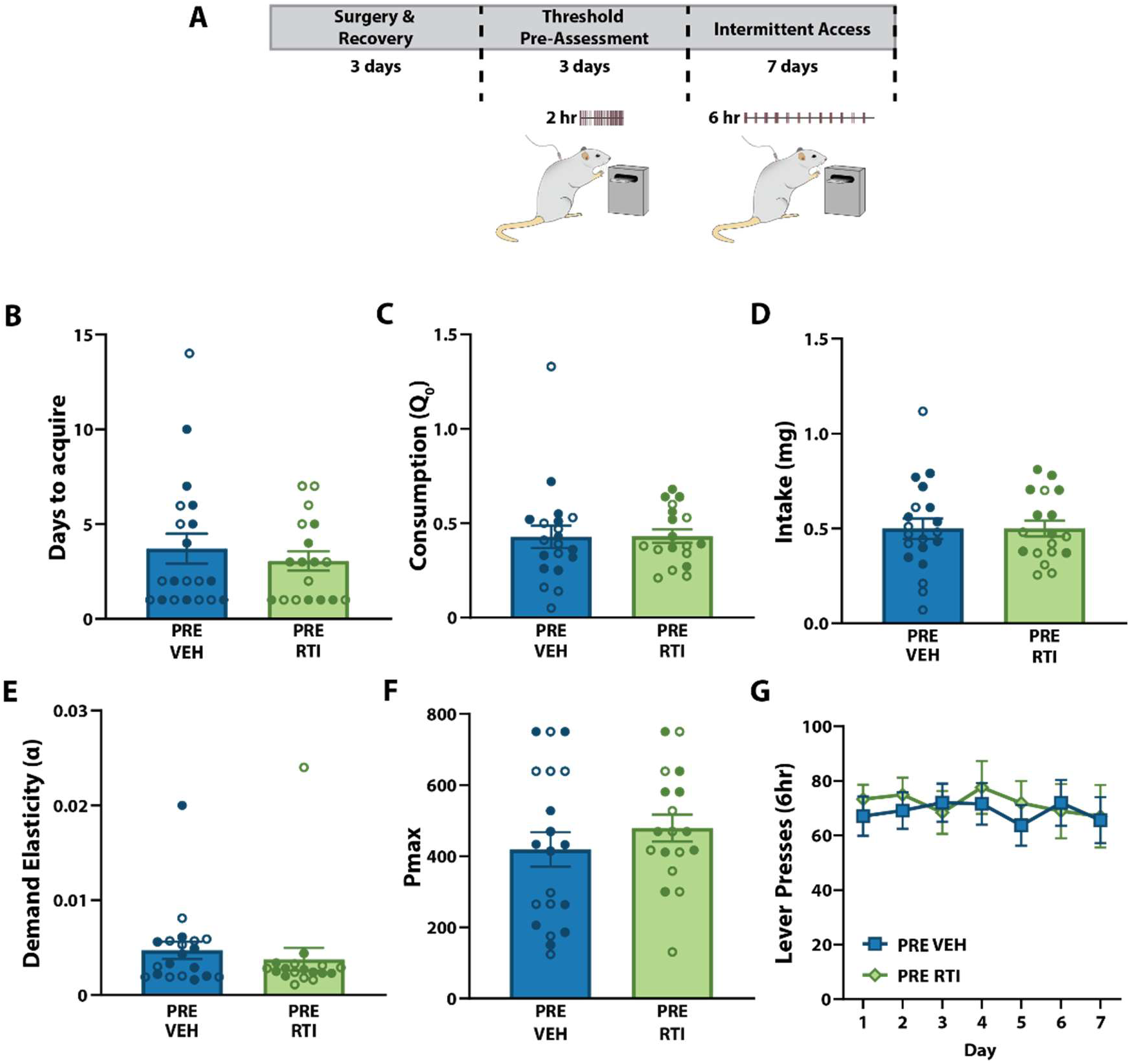
No differences in self-administration prior to vehicle or RTIOX-276 treatment. **A)** Experimental timeline. **B)** Number of days to acquire cocaine self-administration for pre-vehicle (Pre-Veh) and pre-RTIOX-276 (Pre-RTI) groups. **C)** Pre-assessment of consumption (Q0), **D)** intake, **E)** demand elasticity (α), and **F)** Pmax (maximal ‘price’ paid in lever presses/mg cocaine). **G)** Active lever presses during intermittent access (IntA) sessions. Data shown as mean±SEM. ○ females, • males. VEH, n = 20 (10 female & 10 male); RTIOX, n=18 (9 females & 9 males).

Next, we assessed the effects of AD1 treatment with RTIOX-276 on post-assessment measures of consumption after abstinence (**Figure 2A**). A two-way ANOVA with RTIOX-276 treatment (vehicle vs RTIOX-276) as the between subjects variable and threshold timepoint (pre vs post) as the within-subjects variable revealed no effect of RTIOX-276 treatment (*F*_(1,16)_=1.755, *p*=0.3450), timepoint (*F*_(1,16)_=1.755, *p*=0.2039) nor RTIOX-276 treatment x timepoint interaction *(F_(_*_1,16)_=1.405, *p*=0.2531) for consumption (**Figure 2B**). Similarly, a two-way ANOVA with the RTIOX-276 treatment (vehicle vs RTIOX-276) as the between subjects variable and timepoint assessment (pre vs post) as the within-subjects variable revealed no effect of RTIOX-276 treatment *(F_(_*_1,16)_=3.080, *p*=0.0984), timepoint *(F_(_*_1,16)_=1.979, *p*=0.1786) nor RTIOX-276 treatment x timepoint interaction *(F_(_*_1,16)_=1.537, *p*=0.2329) on intake (**Figure 2C**).

Next, we assessed the effects of RTIOX-276 on post-assessment measures of motivation. A two-way ANOVA with the RTIOX-276 treatment (vehicle vs RTIOX-276) as the between subjects variable and threshold timepoint (pre vs post) as the within-subjects variable revealed a significant effect of timepoint (*F*_(1,16)_=6.005, *p*=0.0261) and an RTIOX-276 treatment x timepoint interaction (*F*_(1,16)_=8.417, *p*=0.0104; **Figure 2D**), but no effect of RTIOX-276 treatment (*F*_(1,16)_=0.9542, *p*=0.3432). Bonferroni post-hoc tests indicated that RTIOX-276 significantly increased demand elasticity (α), relative to vehicle treatment. Similarly, a two-way ANOVA with RTIOX-276 treatment (vehicle vs RTIOX-276) as the between subjects variable and threshold timepoint (pre vs post) as the within-subjects variable indicated a significant timepoint effect (*F*_(1,16)_=7.852, *p*=0.0128) and a significant RTIOX-276 treatment x timepoint interaction (*F*_(1,16)_= 8.219, *p*=0.0112; **Figure 2E**), but no effect of RTIOX-276 treatment (*F*_(1,16)_=1.695, *p*=0.2114) on Pmax. Bonferroni tests revealed that RTIOX-276 significantly reduced Pmax relative to vehicle treatment.

**Figure 2:**
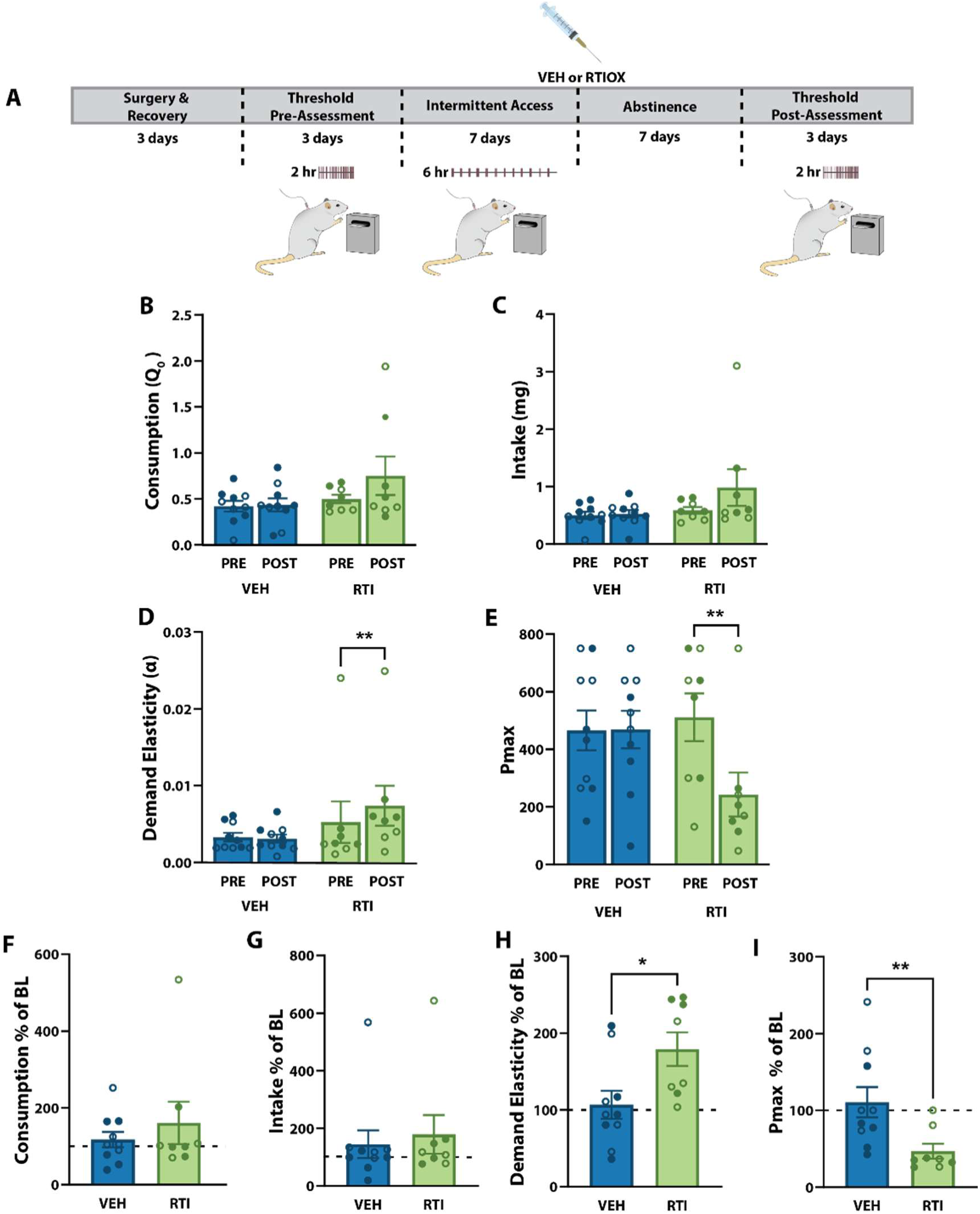
Hcrtr1 blockade early in abstinence reduces motivation for cocaine later in abstinence. **A)** Experimental timeline. **B)** Post-assessment of consumption (Q0), (**C**) intake, (**D**) demand elasticity (α), and **E)** Pmax (maximal “price” paid in lever presses/mg cocaine). Post assessment expressed as a percentage of baseline (BL) of **F)** consumption, **G)** intake, **H)** demand elasticity, and **I)** Pmax. Data shown as mean±SEM. ○ females, • males. VEH, n = 10 (5 female & 5 male); RTIOX, n=8 (4 females & 4 males). (D & E) Two-way ANOVA, Bonferroni; **p<0.01 pre vs post RTIOX, (H & I) Student’s t-tests, *p < 0.05, **p<0.01.

We then determine if post-assessment measures differed when expressed as a percentage of the pre-assessment baseline measures. A Student’s t-tests revealed no significant difference between vehicle and RTIOX treatment for consumption (t_(16)_=0.4227, *p*=0.6782; **Figure 2F**) or intake (t_(16)_=0.8036, *p*=0.2167; **Figure 2G**) when expressed as a percentage of the pre-assessment baselines. By comparison, Student’s t-tests showed that RTIOX-276 significantly increased demand elasticity (t_(16)_=2.574, *p*=0.0102; **Figure 2H**) and decreased Pmax (t_(16)_=2.657, *p*=0.0086; **Figure 2I**) when expressed as a percentage of the pre assessment baselines. Together, these data demonstrate that a single treatment of RTIOX-276 early in abstinence exerts lasting reductions in motivation for cocaine without affecting low effort consumption.

### Hcrtr1 antagonism early in abstinence prevents increases in dopamine uptake following IntA to cocaine

To assess whether a single treatment with RTIOX-276 on the first day of abstinence prevents aberrant dopamine transmission following IntA to cocaine, a separate group of rats was treated with vehicle (n=10) or RTIOX-276 (n=10) on AD1 and baseline dopamine release and uptake as well as dopamine responses to cocaine were assessed in NAc core slices on AD8 using FSCV (**Figure 3A**). Naive rats were used as controls (n=10). A one-way ANOVA with RTIOX-276 treatment as a between subject variable (naive, vehicle, vs RTIOX) showed no effect of RTIOX-276 treatment on dopamine release (*F*_(3,30)_=1.021, *p*=0.3736; **Figure 3B and C**) but there was an effect of RTIOX-276 treatment on dopamine uptake (*F*_(3,30)_=5.064, *p*=0.0136; **Figure 3B and D**). Bonferroni tests revealed a significant increase in dopamine uptake in vehicle-treated rats compared to naive rats and that RTIOX-276 reduced uptake to levels observed in naive rats.

We next examined the dopamine response to cocaine challenge across cumulative concentrations of cocaine (0.3, 1, 3, 10, and 30 µM). A two-way repeated measure ANOVA with RTIOX-276 treatment (vehicle vs RTIOX-276) as the between-subjects variable and cocaine concentration as the within-subjects variable revealed significant effects of cocaine concentration (Greenhouse-Geisser correction; *F*_(2.255,60.87)_=1.235, *p*<0.0001), but no effect of RTIOX-276 treatment *(F*_(2,27)_=1.643, *p*=0.2122) nor a RTIOX-276 treatment x concentration interaction *(F*_(8,108)_=1.235, *p*=0.2857) on dopamine release (**Figure 3F**). Similarly, a two-way repeated measures ANOVA with RTIOX-276 treatment as the between-subjects variable and cocaine concentration as the within-subjects variable revealed a significant effect of cocaine concentration (*F*_(4.108)_=251.0, *p*<0.0001), but no effect of RTIOX-276 treatment (*F*_(2,27)_=1.493, *p*=0.2427; Figure **3G**) nor a RTIOX-276 treatment x cocaine concentration interaction (*F*_(8,108)_=0.5298, *p*=0.8318) on inhibition of dopamine uptake (**Figure 3G**). When considered together, these results indicate that IntA to cocaine significantly increased dopamine uptake rate without affecting dopamine release at baseline or dopamine responses to cocaine. Importantly, RTIOX-276 treatment on AD1 prevented the observed increases in dopamine uptake after IntA to cocaine suggesting the possibility that preventing increases in dopamine uptake may contribute to reductions in motivation for cocaine.

**Figure 3:**
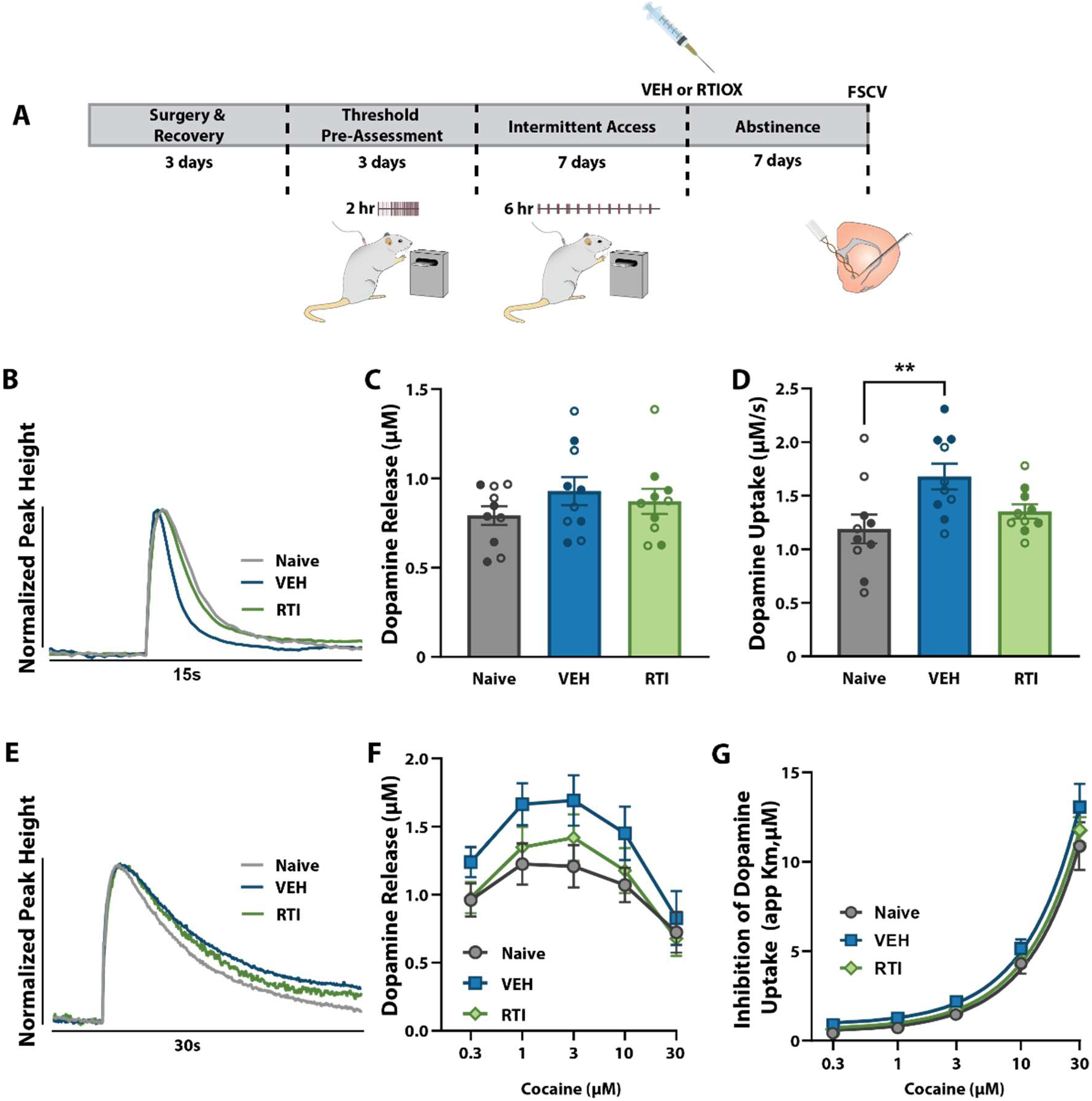
Hcrtr1 blockade normalized aberrant dopamine uptake after IntA to cocaine. **(A)** Experimental timeline. **(B)** Example baseline dopamine traces. Baseline measures of **(C)** dopamine release and **(D)** dopamine uptake for vehicle (Veh) and RTIOX-276 (RTI) treatment. **(E)** Example traces following 30 μM cocaine. **(F)** Cocaine-induced dopamine release and **(G)** DAT sensitivity to cocaine (inhibition of dopamine uptake; app Km). Data shown as mean±SEM. ○ females, • males. Naive, n=10 (5 females & 5 males); VEH, n = 10 (5 female & 5 male); RTIOX, n=10 (5 females & 5 males). One-way.ANOVA, Bonferroni ** *p* < 0.05 Naive vs VEH.

## Discussion

In the present studies we assessed the effects of Hcrtr1 blockade on consumption and motivation for cocaine and dopamine transmission following abstinence from IntA to cocaine. We found that treatment with a Hcrtr1 antagonist on the first day of abstinence produced lasting reductions in motivation for cocaine after 7 days of abstinence but had little effect on low effort consumption. Additionally, we found that IntA cocaine self-administration enhanced the efficiency of dopamine uptake after 7 days of abstinence and that Hcrtr1 antagonism prevented this increase.

### Blockade of Hcrtr1 early in abstinence exerts lasting reductions in motivation for cocaine

In the present study, we hypothesized that abstinence from IntA would intensify motivation for cocaine as previously reported with a similar within-session threshold procedure (Calipari et al., 2015), and because abstinence following cocaine use engenders incubation of cocaine seeking (Grimm et al., 2001; Grimm et al., 2003; Lu et al., 2004; Wolf, 2016; Alonso et al., 2022; Clark et al., 2024). Surprisingly, we found that 7-days of abstinence from IntA did not increase measures of consumption or motivation for cocaine during the within-session threshold sessions (i.e., in vehicle-treated rats). Rather, we observed maintained consumption, intake, demand elasticity, and Pmax levels between pre- and post-assessment measures. These results differ from previous work, where as few as 3 days of IntA to cocaine followed by 7 days of abstinence reduced consumption and increased motivation for cocaine (Calipari et al., 2015). The reasons for these differences are unclear – yet when comparing our pre-assessment measures to the previous report it is evident that our rats expressed greater Pmax prior to IntA to cocaine. Given that Pmax responding has an inherent ceiling, the high Pmax levels observed during our pre-assessment sessions limits how much higher motivation for cocaine could be during the post-assessment sessions (**Supplemental Figure 1**).

Ideal interventions for relapse to cocaine use would occur early in abstinence to curtail motivation and craving, therefore reducing likelihood or severity of relapse. In the present studies, we hypothesized that blockade of Hcrtr1 early in abstinence would prevent motivation for cocaine following abstinence. Indeed, recent work from our laboratory indicates that Hcrtr1 blockade on the first day of abstinence from IntA to cocaine prevents incubation of cocaine seeking later in abstinence (Clark et al., 2024). Consistent with these findings, in the current studies we found that RTIOX-276 on the first day of abstinence increased demand elasticity and decreased Pmax – indicating a reduction in motivation for cocaine. Therefore, despite failing to observe increased motivation after abstinence from IntA to cocaine in vehicle-treated rats, we nevertheless observed a considerable decrease in these measures following RITOX-276 – indicating that HcrtR1 blockade exerts lasting reductions in motivation for cocaine.

### Hcrtr1 blockade normalizes dopamine uptake following abstinence from IntA to cocaine

The present studies revealed that abstinence from IntA to cocaine increased baseline dopamine uptake rate in vehicle-treated rats. This elevation in dopamine uptake is similar to what has been reported in early (Calipari et al., 2015; Clark et al., 2024) and later (Alonso et al., 2022) abstinence. As predicted, a single dose of RTIOX-276 on the first day of abstinence normalized increases in dopamine uptake observed following abstinence to IntA to cocaine, which is similar to what we observed recently in rats that displayed incubation of cocaine seeking (Clark et al., 2024).

In the current studies, however, we did not observe changes in dopamine release following IntA and abstinence which differs from some prior observations (Calipari et al., 2013) including our recent studies (Clark et al., 2024). Nonetheless, other studies examining the effects of IntA and abstinence have also failed to report changes in dopamine release (Calipari et al., 2014; Alonso et al., 2022), suggesting that abstinence from cocaine may not consistently affect dopamine release and related mechanisms.

### Blockade of Hcrtr1 during abstinence did not alter the dopamine response to cocaine

Based on previous reports, we hypothesized that IntA exposure to cocaine would increase DAT sensitivity to cocaine. Although we observed a non-significant increase in dopamine release after cocaine in vehicle-treated rats following abstinence from IntA to cocaine, we did not see changes in DAT sensitivity to cocaine. These findings are in conflict with prior studies indicating IntA to cocaine increases DAT sensitivity to cocaine following 1 day or 7 days of abstinence (Calipari et al., 2015). Moreover, in our recent studies we also observed greater DAT sensitivity to cocaine at early (Clark et al., 2024) or later (Alonso et al., 2022) abstinence timepoints following IntA to cocaine. Interestingly, after 7 days of abstinence only rats that displayed incubation of cocaine seeking showed increases in dopamine responses to cocaine, while those that did not incubate had similar dopamine transmission patterns as cocaine-naive rats (Clark et al., 2024). Given the experimental design in the current studies, which used separate rats for examining post abstinence behavioral measures than those used for FSCV measures, we are unable to link potential differences in cocaine seeking to FSCV changes in the same rats. As such, we cannot assess if individual differences in exaggerated motivation for cocaine could explain a lack of effects on dopamine responses to cocaine observed herein.

### Lasting effects of single treatment with Hcrtr1 antagonists

The vast majority of studies have examined the acute, on-board pharmacological effects of Hcrt antagonists. In the context of cocaine-associated behavior, on-board Hcrtr1 blockade reduces high effort motivation for cocaine (Borgland et al., 2009; España et al., 2010; Brodnik et al., 2015; Prince et al., 2015; Levy et al., 2017; Pantazis et al., 2022), as well as cocaine intake under restricted access schedules (España et al., 2010). These acute actions of HcrtR1 antagonists are also observed for changes in dopamine transmission including reductions in baseline release and uptake as well as attenuation of dopamine responses to cocaine (España et al., 2010; Prince et al., 2015; Levy et al., 2017). However, a limited set of reports have begun to reveal effects HcrtR1 antagonists that are long lasting – extending beyond on-board, pharmacological effects (Ishii et al., 2005; Winrow et al., 2010; Brodnik et al., 2020; Mohammadkhani et al., 2020). For instance, Hcrtr1 antagonism exerts lasting effects on behavior and dopamine transmission with reduced cocaine self-administration observed after 21 hr of treatment and attenuated dopamine uptake and DAT sensitivity to cocaine after 24 hr – both timepoints which are beyond the pharmacological availability of the compound (Brodnik et al., 2020). Further, we recently reported that a single treatment of RTIOX-276 on the first day of abstinence prevents incubation of cocaine seeking and normalizes dopamine transmission following abstinence from IntA to cocaine (Clark et al., 2024). These latter observations are consistent with our present findings, suggesting that Hcrtr1 blockade produces lasting effects that impact dopamine transmission and motivation for cocaine.

## Summary

In the present studies, we examined the lasting effects of Hcrtr1 blockade on motivation for cocaine and changes in dopamine transmission after 7 days of abstinence from IntA to cocaine. We observed that a single treatment with the Hcrtr1 antagonist, RTIOX-276, on the first day of abstinence reduced motivation without affecting low effort consumption of cocaine. We also found that IntA to cocaine followed by 7 days of abstinence enhanced dopamine uptake and this effect was prevented by RTIOX-276 treatment on the first day of abstinence. We did not observe differences in DAT sensitivity to the effects of cocaine following IntA and abstinence, nor did RTIOX-276 have significant effects on these measures. Together these observations reveal that Hcrtr1 blockade produces lasting reductions in motivation when delivered early in abstinence to prevent motivation for cocaine which may be linked to attenuation of dopamine uptake.

## Acknowledgements

We thank the NIDA drug supply program for donating the cocaine hydrochloride. This work was supported by the National Institute on Drug Abuse grant R01DA039100 (RAE) and National Science Foundation GRFP Grant No. 2041772 (SBS).

## Author Contributions

SBS conducted the self-administration and FSCV data collection, analyzed and interpreted the data, wrote the original draft, and edited the content. JKS, conceptualized and designed the studies, conducted the self-administration and FSCV data collection. IPA and EMB conducted the self-administration and FSCV data collection. RAE, conceptualized and designed the studies, analyzed and interpreted the data, edited the original and subsequent drafts.

