## Supplemental Materials for "Hypocretin receptor 1 blockade early in abstinence reduces future demand for cocaine"

**Supplemental Table 1. RTIOX-276 Sex differences analyses.**

| Measures | Dependent Variable | Results | Test |
|---|---|---|---|
| Days to Acquire | Sex<br>Group (Vehicle, RTIOX-276)<br>Group x Sex | F(1,34)=0.2227, p=0.6400<br>F(1,34)=0.3287, p=0.5702<br>F (1,34)=2.446, p=0.1271 | Two-way ANOVA |
| Pre-Assessment Consumption | Sex<br>Group (Vehicle, RTIOX-276)<br>Group x Sex | F(1,34)=0.8240, p=0.3704<br>F(1,34)=0.001966, p=0.9649<br>F(1,34)=1.011, p=0.3217 | Two-way ANOVA |
| Pre-Assessment Intake | Sex<br>Group (Vehicle, RTIOX-276)<br>Group x Sex | F(1,34)=0.0.09908, p=0.7549<br>F(1,34)=0.4768, p=0.4946<br>F(1,34)=0.2043, p=0.6542 | Two-way ANOVA |
| Pre-Assessment Deman Elasticity | Sex<br>Group (Vehicle, RTIOX-276)<br>Group x Sex | F(1,34)=0.0536, p=0.8183<br>F(1,34)=0.007337, p=0.9322<br>F(1,34)=2.201, p=0.1471 | Two-way ANOVA |
| Pre-Assessment Pmax | Sex<br>Group (Vehicle, RTIOX-276)<br>Group x Sex | F(1,34)=0.2546, p=0.6171<br>F(1,34)=0.8867, p=0.3530<br>F(1,34)=0.3196, p=0.5756 | Two-way ANOVA |
| Lever Presses IntA | Day (1-7)<br>Group (Vehicle, RTIOX-276)<br>Sex<br>Day x Sex<br>Group x Sex<br>Group x Day<br>Day x Group x Sex | F(6,204)=1.009, p=0.4205<br>F(1,34)=0.001697, p=0.9674<br>F(1,34)=7.659 p=0.0091**<br>F(6,204)=3.573, p=0.0022**<br>F(1,34)=0.4418, p=0.5108<br>F(6,204)=0.8319 p=0.5464<br>F(6,204)=0.6009, p=0.7294 | Three-way mixed Design ANOVA |
| Post-Assessment Consumption | Timepoint (pre, post)<br>Sex<br>Group (Naive, Vehicle, RTIOX-276)<br>Timepoint x Sex<br>Timepoint x Group<br>Group x Sex<br>Timepoint x Sex x Group | F(1,14)=1.588, p=0.2282<br>F(1,14)=0.1925, p=0.6675<br>F(1,14)=2.635, p=0.1269<br>F(1,14)=0.3956, p=0.5395<br>F(1,14)=1.272, p=0.2783<br>F(1,14)=0.02923, p=0.8667<br>F(1,14)=0.1326, p=0.7212 | Three-way ANOVA |
| Post-Assessment Intake | Timepoint (pre, post)<br>Sex<br>Group (Naive, Vehicle, RTIOX-276)<br>Timepoint x Sex<br>Timepoint x Group<br>Group x Sex<br>Timepoint x Sex x Group | F(1,14)=1.921, p=0.0.1875<br>F(1,14)=0.8852, p=0.02163<br>F(1,14)=0.1193, p=2.752<br>F(1,14)=0.2513, p=1.432<br>F(1,14)=0.2421, p=1.492<br>F(1,14)=0.6212, p=0.2553<br>F(1,14)=0.6666, p=0.1936 | Three-way ANOVA |
| Post-Assessment Demand Elasticity | Timepoint (pre, post)<br>Sex<br>Group (Naive, Vehicle, RTIOX-276)<br>Timepoint x Sex<br>Timepoint x Group<br>Group x Sex<br>Timepoint x Sex x Group | F(1,14)=6.586, p=0.0224*<br>F(1,14)=0.1349, p=0.7189<br>F(1,14)=1.572, p=0.2305<br>F(1,14)=1.527, p=0.2369<br>F(1,14)=9.232, p=0.0089*<br>F(1,14)=0.7635, p=0.3970<br>F(1,14)=2.404, p=0.1434 | Three-way ANOVA |
| Post-Assessment Pmax | Timepoint (pre, post)<br>Sex<br>Group (Naive, Vehicle, RTIOX-276)<br>Timepoint x Sex | F(1,14)=9.520, p=0.0081*<br>F(1,14)=1.110, p= 0.3099<br>F(1,14)=0.9792, p=0.3392<br>F(1,14)=5.388, p=0.0359* | Three-way ANOVA |

| | | | |
|---|---|---|---|
| | Timepoint x Group<br>Group x Sex<br>Timepoint x Sex x Group | F(1,14)=9.966, p=0.0070*<br>F(1,14)=1.041, p=0.3250<br>F(1,14)=0.1354, p=0.7184 | |
| Baseline Dopamine Release | Sex<br>Group (Naive, Vehicle, RTIOX-276)<br>Group x Sex | F(1,24)=0.5007, p=0.4860<br>F(2,24)=0.9276, p=0.4092<br>F(2,24)=0.01176, p=0.9883 | Two-way ANOVA |
| Baseline Dopamine Uptake | Sex<br>Group (Naive, Vehicle, RTIOX-276)<br>Group x Sex | F(1,24)=0.3614, p=0.8508<br>F(2,24)=4.941, p=0.0160*<br>F(2,24)=1.153, p=0.3327 | Two-way ANOVA |
| Effect of Cocaine on Dopamine Release | Concentration (0.3 - 30μM)<br>Sex<br>Group (Naive, Vehicle, RTIOX-276)<br>Sex x Group<br>Concentration x Sex<br>Concentration x Group<br>Concentration x Group x Sex | F(2.136,51.259)=51.196, p<0.001<br>F(1,24)=0.967, p=0.335<br>F(2,24)=1.443, p =0.256<br>F(2,24)=0.075, p=0.928<br>F(2.136,51.259)=0.936, p=0.404<br>F(4.272,51.259)=1.259, p=0.297<br>F(4.272,51.259)=1.258, p=0.298 | Three-way Mixed Design ANOVA |
| Effect of Cocaine on Inhibition of Dopamine uptake | Concentration (0.3 - 30μM)<br>Sex<br>Group (Naive, Vehicle, RTIOX-276)<br>Sex x Group<br>Concentration x Sex<br>Concentration x Group<br>Concentration x Group x Sex | F(1.095,26.278)=271.714, p<0.001<br>F(1,24)=1.288, p=0.268<br>F(2,24)=1.584, p =0.226<br>F(2,24)=0.385, p=0.994<br>F(1.095,26.278)=0.506, p=0.5<br>F(2.190,26.278)=0.577, p=0.583<br>F(2.190,26.278)=1.096, p=0.354 | Three-way Mixed Design ANOVA |

**Discussion**

Our current findings indicate no sex differences in days to acquire, pre-assessments for consumption, intake, demand elasticity or Pmax. These findings are different from previous reports utilizing similar within session threshold experiments that indicate that females have lower demand elasticity (greater motivation) compared to males (Kohtz et al., 2022). However, it is important to note that in our experiments we observed relatively low demand elasticity (high motivation) across both sexes to levels comparable to those reported for only females. We observed significant sex differences in the amount of lever presses during intermittent access (IntA) to cocaine, in which females press more than males similar to previous findings that females are more motivated to self-administer cocaine (Roberts et al., 1987; Roberts et al., 1989; Carroll et al., 2002; Lynch, 2006). Our current findings indicate no overall sex differences or an interaction between sex and RTIOX-276 treatment in post-assessments of for consumption, intake, demand elasticity or Pmax, indicating that the behavioral effects of RTIOX-276 did not differ across sex. Further, we did not observe overall sex differences or an interaction between sex and RTIOX-276 treatment for voltammetry measures of baseline or cocaine-induced dopamine release or uptake, again indicating that the dopamine effects of RTIOX-276 did not differ across sex. These findings differ from previous reports indicating that females have greater dopamine release and uptake within the striatum (Walker et al., 2000; Walker et al., 2006; Calipari et al., 2017).

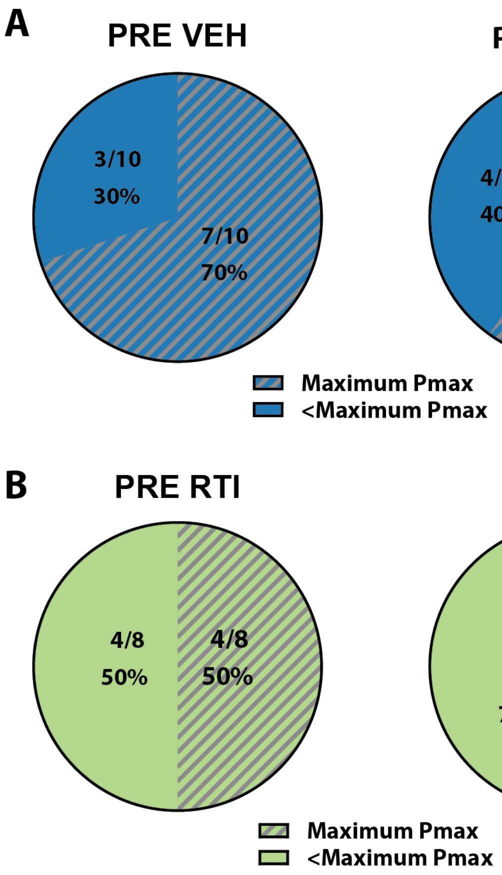


**Supplemental Figure 1: Within-session threshold ceiling effect for Pmax.** During within-session threshold sessions rats can achieve a maximum Pmax of 750. (**A**) Proportion of vehicle-treated (VEH) rats that reached the maximum Pmax on at least one session during pre- or post-assessments. (**B**) Proportion of RTIOX-276-treated rats (RTI) that reached the maximum Pmax on at least one session during pre- or post-assessments.
